# Monocotyledonous vascular bundles characterized by image analysis and pattern recognition

**DOI:** 10.1101/494039

**Authors:** Juliana V. da Silva, Cesar H. Comin, Luciano da F. Costa

## Abstract

Recent advances in image analysis and pattern recognition have paved the way to several developments in plant science. In the present work, we report the comparative study, by using the aforementioned approaches, of vascular bundles of *Dracaena marginata*. More specifically, we used 33 measurements related to shape, density and regularity of imaged cross-sections of the stem. By using individual, pairwise and PCA projections of the adopted measurements, we were able to find the combinations of measurements leading to the best separation between the considered tissues. In particular, the best separation was obtained for entropy taken at a particular spatial scale combined with the equivalent diameter. The reported developments open several perspectives for applications in content-based retrieval, diagnosis, and species identification.

## I. INTRODUCTION

An organism can be defined as a group of organs working together to perform one or more functions (Raven et al., 2014). Each organ is constituted by tissues, which are ensembles of similar cells. In plants, the study of tissues can be divided into two types: meristematic and permanent. Meristematic tissues, also called embryonic, are composed of cells in constant division. These tissues are of vital importance for plants growth in length and thickness, being also responsible for the derivation of all other tissues. Examples of this type of tissue are: protoderm, procambium, cambium and felogen. The permanent tissues are composed by differentiated cells that have permanent shape, size and a function. There are two types of permanent tissues: simple permanent tissues (for example: parenchyma, collenchyma, sclerenchyma, epidermis) and complex permanent tissues (for example: xylem and phloem) (Raven et al., 2014)

Xylem and phloem are plant tissues important for transporting fluid and nutrients internally. The former provides the main way to transport water and mineral salts (raw sap) from roots into leaves, whereas the latter is a tissue conducting elaborated sap (organic substance, such as sugars, in solution) from leaves into roots. The xylem and phloem can be primary or secondary depending on the respective embryonic origin: procambium or cambium, respectively (Raven et al., 2014). Both xylem and phloem are known as plant vascular tissues and, although they have no function in plant growth, are present when growth occurs in length and thickness in the plant.

The primary vascular tissue, composed by primary xylem and phloem, is involved in the longitudinal growth, known as *primary growth*. In dicotyledonous, this tissue is organized as concentric hollow cylinders in which the internal cylinder is delimited by procambium and the external is delimited by protoderm. The development of xylem is endogenous in relation to procambium, whereas the development of phloem is exogenous. On the other hand, in most monocots, the primary vascular bundles are produced by procambium and, subsequently, the pericycle participates in the formation of these bundles (Menezes et al., 2005; Lima and Menezes, 2009; Cattai and Menezes, 2010). These vascular bundles can occur in five different types: collateral closed, collateral open, bicolateral, concentric periphloematic (also know as amphicribral) and concentric perixylematic (also know as amphivasal).

The secondary vascular tissue is composed by secondary xylem and phloem and is related to the diametric growth, known as *secondary growth*. In general, in dicotyledonous, the procambium will originate the cambium which, in turn, will develop secundary xylem and phloem. Usually, monocots do not have secondary growth. However, some species of this class, such as *Dracaena draco, Dracaena marginata, Yucca aloifolia, Yucca brevifolia, Cordyline fruticosa*, among others (Tomlinson and Zimmermann, 1969), have secondary growth and, consequently, secondary vascular bundles.

The vascular system of monocots with thickness growth is less known as a consequence of its complexity. It has been studied since the 19th century to the present day in works such as: Mirbel (1809), de Mirbel (1843), von Mohl (1849), Metcalfe et al. (1960), Tomlinson (1964), Tomlinson and Zimmermann (1969), Menezes et al. (2005), Lima and Menezes (2009) and Cattai and Menezes (2010). The work of the Tomlinson and Zimmermann (1969) shows that some studies believe that the secondary thickening meristem originates from pericycle divisions, therefore being longitudinally continuous and having histological and functional similarity. In these cases, the pericycle will give rise to the secondary thickening meristem, known as monocot cambium Carlquist (2012), which will produce the secundary vascular bundles. One of the works cited by Tomlinson and Zimmermann (1969) is Hausmann (1908), which shows that the distinction between primary and secondary meristems is complex and consequently the characterization of primary and secondary vascular tissues is also difficult.

Among the group of monocotyledons with secondary growth, the genus *Dracaena* has been studied for a long time, but there are still gaps to be filled about the characterization of the vascular tissues of these plants. According to the work of Carlquist (2012) these studies were limited by the knowledge and, mainly, by the techniques available at the time. That is, most of the research on the vascular system of monocotyledons considers histology based only on optical microscopy to recognize, differentiate and characterize tissues (Metcalfe et al., 1960; Tomlinson, 1964; Tomlinson and Zimmermann, 1969).

The characterization of plant tissues consisted in identifying the histological visual characteristics that made it a unique tissue, that is, it was a relatively subjective technique based on human cognition. Currently, the continuing advances in image analysis concepts and methods, allied to the ever improving pattern recognition approaches, have paved the way to obtaining increasingly more comprehensive and accurate characterizations of fruit plant tissue as described in Ramos and et al. (2004), Mayor et al. (2005), Mayor et al. (2008) and Sanyal et al. (2008) in which it is possible to study the effect of dehydration of a tissue in several organisms, and thus make inferences about the quality of the fruit.

The search for microhistological descriptors through the analysis of plant images has contributed to identifying species (Rosito and Marchesan, 2003) and to understanding the tissue architecture during its development or when affected by diseases (Sánchez-Gutiérrez et al., 2016). Within this context, the present work set out to harness the benefits of digital image analysis and pattern recognition in order to better understand plant tissues, especially in the sense of achieving more quantitative and objective results, and generating new interpretations in the characterization of monotiledoneous vascular tissues with secondary growth. In particular, we aim at finding descriptors that contribute to effective differentiation between primary and secondary vascular bundles and that have biological relevance aiding the understanding of the architecture of these tissues.

As observed above, among the group of monocotyledons with secondary growth, the genus *Dracaena* is the most studied. However, the choice of a particular species is inherently limited by geographical availability. So, *Dracaena marginata* was chosen in this work because of two main reasons: (i) This species is easily cultivated in tropical and subtropical climates; and (ii) It has both primary and secondary vascular bundles of the amphivasal type, allowing a more controlled investigation.

So, the present study aims at differentiating the primary from the secondary vascular bundles of *Dracaena marginata* in terms of a systematic application of digital image analysis and pattern recognition. We analyzed 20 images of primary and secondary vascular bundles of *Dracaena marginata* in order to try to characterize these tissues and to search for attributes that were capable of not only segregating them computationally, but also that had biological significance. Images were manually segmented using *Paint.net* software. We extracted 33 attributes divided into 3 classes: shape measurements, density measurements and structural regularity measurements. These attributes were then analyzed by 3 different methodologies: individual measurement analysis; pairwise measurement analysis and principal component analysis.

The methodology that best addressed the objectives proposed in this study was the pairwise measurement analysis. Respective results showed that the best separation of the tissues was obtained when the measurement entropy with sigma 30 was combined with the average equivalent diameter. We show that these attributes have significant biological relevance and are therefore potential classifiers to distinguish primary and secondary amphivasals vascular bundles. The obtained results pave the way to several applications such as species identification using vascular bundles, as well as diagnosis of environment induced abnormalities and developmental studies. Therefore, the methodology applied in this work can be extended to other plant species, as well as other types of vascular bundles.

## II. RESULTS

### A. Hystologic Interpretation

Monocotyledons that show secondary growth may have vascular bundles of two types: concentric (amphivasal or amphicribral) or collateral (open or closed) (Tomlinson and Zimmermann, 1969; Cattai and Menezes, 2010; Jura-Morawiec, 2017). The species *Dracaena marginata* is characterized by having primary and secondary vascular bundles of the amphivasal type (Jura-Morawiec, 2015).

Figure 1 illustrates images of the transverse histological sections of the stem of the *Dracaena marginata*. In the figure, we can observe that the primary (see Figure 1 A) and secondary (see Figure 1 B) vascular bundles are of the amphivasal type. These vascular bundles are characterized by having the phloem surrounded by the xylem. That is, the analysis of the transverse sections of the stem of *Dracaena marginata*, as done in this work, are consistent with the literature.

**FIG. 1:**
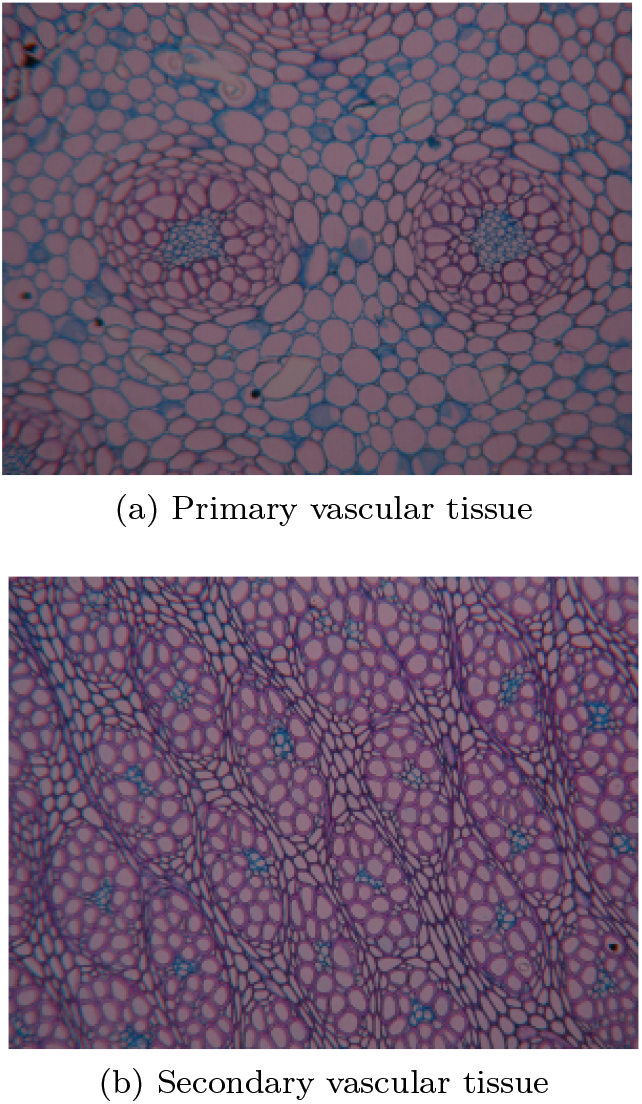
Examples of tissues found in *Dracaena marginata*. Tissues tags in blue are phloem and tissues tags in purple are xylem.

A total of 20 images containing primary and secondary vascular bundles of the stem of *Dracaena marginata* B. was analyzed. A set of 33 morphological, density and regularity measurements (described in Section IV B) was extracted from each image, which was then studied using three approaches: i) individual analysis; ii) pairwise analysis and iii) PCA projection. These approaches are described in detail in Section IV C. The aim was to search for the attributes that could segregate well the primary and secondary vascular bundles of monocotyledon with secondary growth. The results obtained in each approach will be described below.

### B. Individual Measurement Analysis

In order to infer how much a measurement can individually characterize and segregate the vascular tissues from *Dracaena marginata*, graphs based on the probability density function estimation of each measurement were plotted. According to Bayesian inference, the smaller the overlap region between two curves, the better the groups will be separated and, consequently, the classes will be better defined. Graphs with the smallest overlapping region between the probability density functions are shown in Figure 2.

**FIG. 2:**
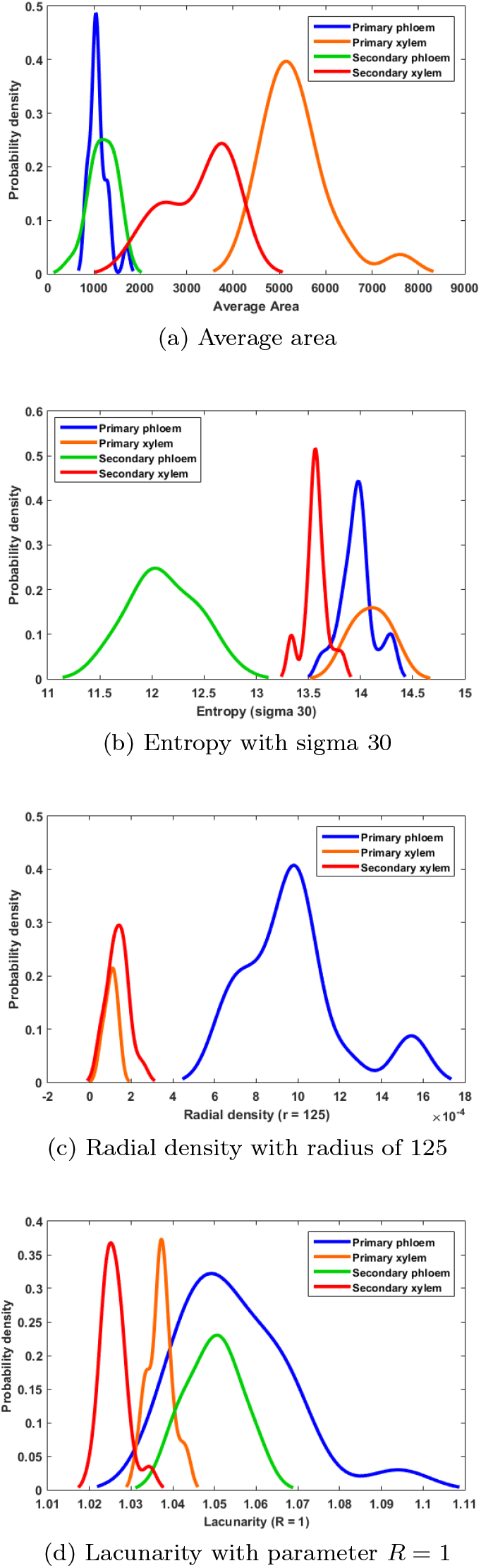
Graphs showing the measurements yielding the best separation between the tissues analyzed.

Figure 2 A, B, C, and D presents the plots obtained for the individual measurements, considering all images. They show, respectively, the density functions of average area, entropy with sigma 30, radial density with radius of 125 and lacunarity with parameter *R* =1.

Analyzing Figure 2 A it is observed that the xylem, both primary and secondary, have density distribution curves of average cell area that have with little overlap between them. Whereas, when evaluating the phloem curves, primary and secondary, there is a great overlap between them. As a consequence, we can infer that the mean cell area measurement was able to separate the primary and secondary xylems from each other better than from the primary and secondary phloem. Another interesting observation about Figure 2 A is that the primary and secondary xylem density curves are shifted towards a larger cell area, whereas the curves of the primary and secondary phloem are centered in the region of the smallest cell area.

Figure 2 B shows the entropy attribute densities. The primary tissues, both phloem and xylem, have density distribution curves very close to the center point and with relatively large dispersions, so they can not be well differentiated by entropy 30. Secondary tissues resulted well separated regarding this measurement. The entropy of an image can be understood as a quantification of randomness. Therefore, the greater the value of this measure, the more irregular the analyzed image will be. When we analyze Figure 2 B we can infer that the tissue that presents the least entropy and greatest regularity, is the secondary phloem. However, its density distribution curve is markedly elongated, indicating that the entropy of this tissue varies substantially among the considered images.

Figure 2 C presents the radial density results for radius equal to 125, from which we observe that there was a significant separation of the primary phloem, which does not present overlap with none of the other tissues. There is no secondary phloem curve. This can be explained by the fact that in this type of vascular bundle, the phloem is enveloped by the xylem. The secondary tissue is smaller than the primary tissue and, therefore, the radius of 125 was not able to measure the internal tissue, that is, the secondary phloem. In this way, we can deduce that with the radius of 125 it was possible to map the primary phloem, but this ray was unsatisfactory to segregate the other tissues.

Another measurement that led to tissue separation was the lacunarity (see Figure 2 D). More specifically, it promoted the separation of primary and secondary xylems one another, and from the others. The behavior of the curves shows that the primary xylem lacunarity is slightly higher than for the secondary xylem. However, this measurement was unable to separate the primary and secondary phloem from each other as evidenced by the large region of overlap between the density function curves.

As shown and discussed above, although some measures have allowed considerable separation between the tissues, none of them was able to completely separate all the tissues analyzed. It is therefore interesting to consider a pairwise combination of measurements, which is developed in the following section.

### C. Pairwise Measurement Analysis

In order to analyze the degree of correlation between the considered measurements, the Pearson’s correlation coefficient between each measurement pair was calculated. The results were organized into two figures: 3 and 4. Figure 3 shows the Pearson correlation matrix. It illustrates, by means of a color scale, how much two measures are pairwise correlated so that the darker the color intensity, the greater the correlation between measurements. The negative values indicates negative correlation. In this figure, two regions of darker shades are observed: (i) the first one, at the top of the matrix, with measurements related to shape; and (ii) a second region, more to the center, involving density related measurements. In order to have a better visualization of the correlations, a network was also plotted as shown in Figure 4. Each node corresponds to one of the measures being studied. Edges are shown whenever −0.9 > *p* and 0.9 < *p*.

**FIG. 3:**
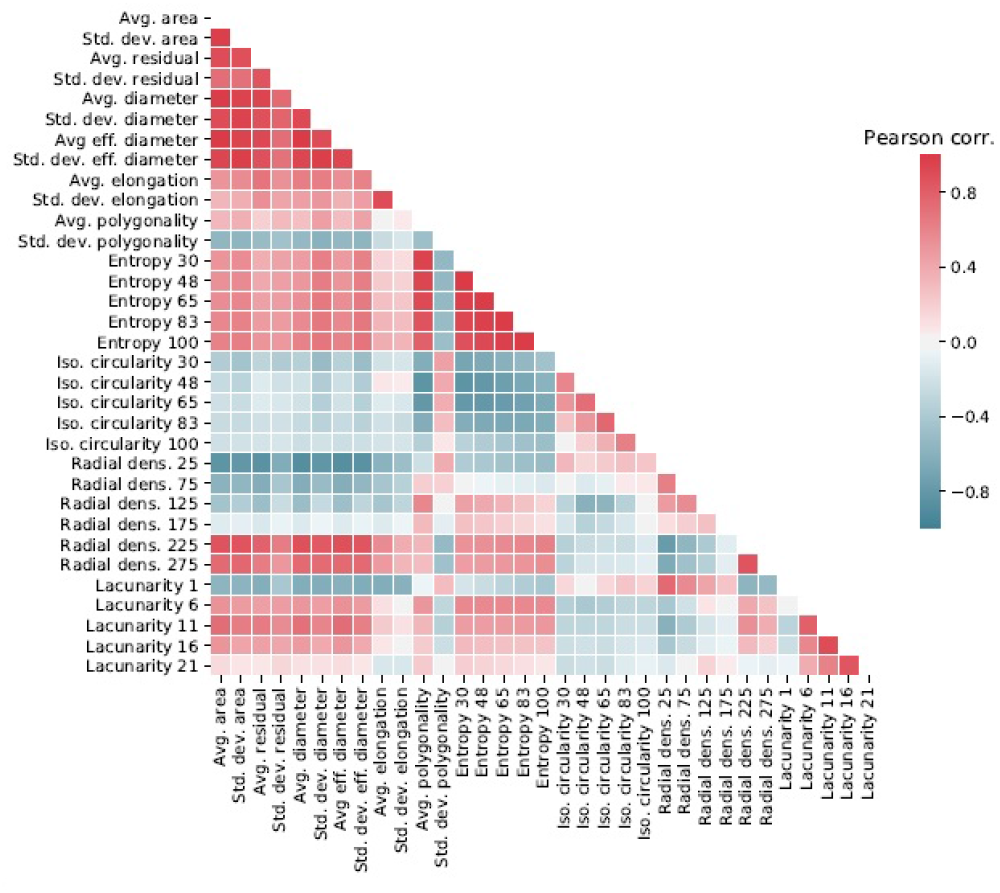
Pearson correlation matrix obtained for pairwise combinations of the adopted features.

**FIG. 4:**
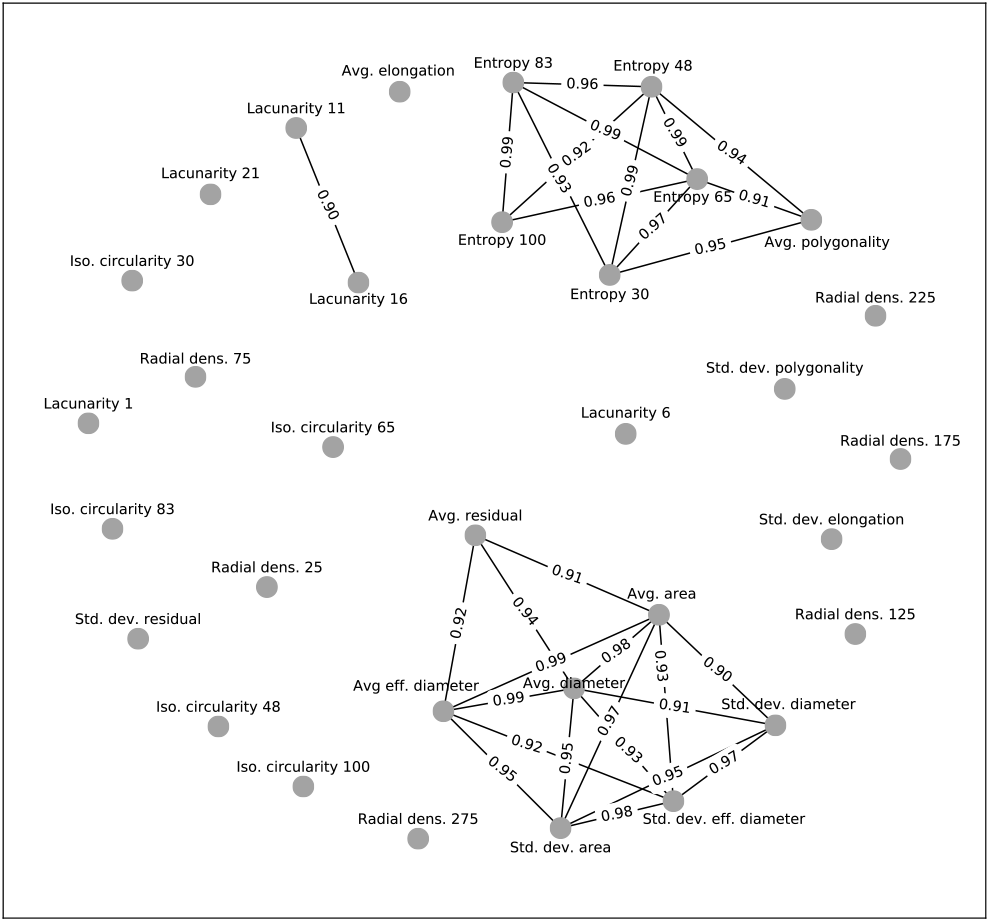
Network with Pearson correlation.

Through the analysis of Figure 4, we have that the measurements concerning the cellular shape have a high correlation one another. On the other hand, out of the measures related to density, only the entropy-related features are correlated one another and with the average polygonality. It is also noted that there is a correlation between lacunarity measures 11 and 16.

After selecting the pairs of attributes with −0.9 < *p* < 0.9, the scatterplots of these measurements were plotted and analysed. A total of 545 graphs were obtained and had to be filtered for analysis so as to select the cases exhibiting better separation between the tissues. In this way, only 5 graphs with largest ratios of inter and intracluster distances were selected, as given in Table I. These graphs are shown in Figure 5 which illustrates the graphs resulting from the combination of: A. Entropy 30 x Average equivalent diameter; B. Entropy 30 x Average area; C. Entropy 30 x Average diameter; D. Entropy 48 x Average equivalent diameter and E. Entropy 48 x Average area.

**TABLE I:** Interclusters and intraclusters average distance and relation (r) between distances for the five selected graphs.

| Plots | Inter | Intra | $r$ |
| --- | --- | --- | --- |
| Entropy 30 x Avg. Eff. Diameter | 60,77 | 11,82 | 5,14 |
| Entropy 30 x Avg. Area | 58,53 | 11,52 | 5,08 |
| Entropy 30 x Avg. Diameter | 63,72 | 12,60 | 5,06 |
| Entropy 48 x Avg. Eff. Diameter | 59,50 | 11,79 | 5,05 |
| Entropy 48 x Avg. Area | 57,24 | 11,54 | 4,96 |

**FIG. 5:**
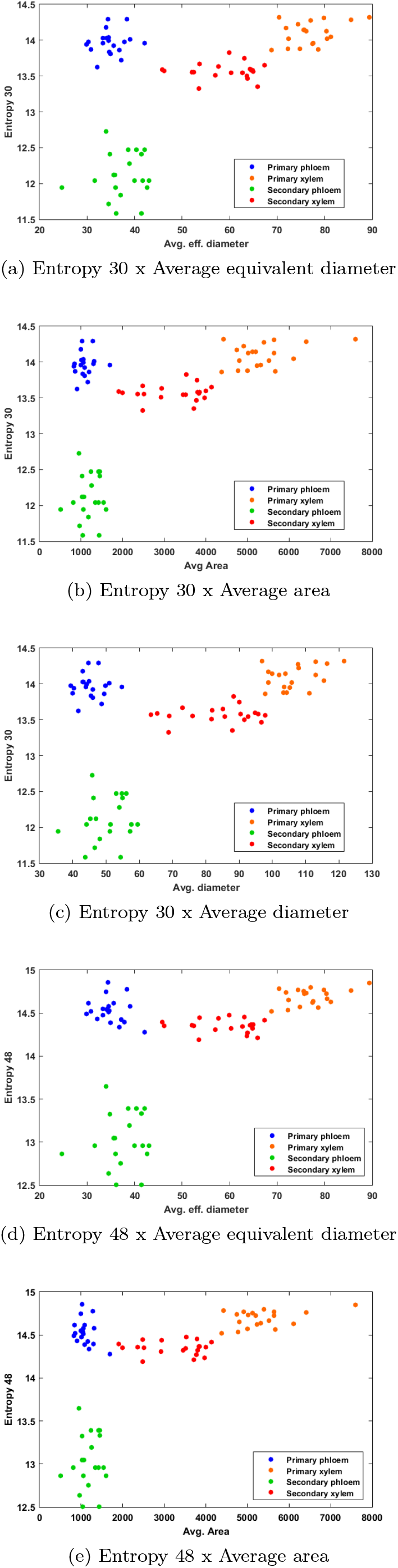
Scatterplots obtained by selecting pairs of measurements exhibiting the largest separations between the categories.

By analysing Figure 5, we observe that the entropy is present in all the selected cases, and in two of them we have the entropy 48 and in the others the entropy 30. The Pearson coefficient between these two measurements is 0.99, as evidenced in Figure 4. The latter attribute had already presented good results in the individual analysis, as it was able to completely segregate the secondary phloem. However, the other tissues had an overlap area.

Among the measurements referring to the cellular shape, the graphs selected were those related to the average equivalent diameter, average diameter and average cell area. The attribute average area, which yielded good results when used individually, appears in two plots. In that individual case, due to the characteristics of the tissues, it was possible to segregate between phloem and xylem, but there were overlaps between the primary and secondary tissues. Another attribute that was also selected twice is the average equivalent diameter. This measurement is based on the cell area. For this reason, average area and average equivalent diameter are strongly correlated as can be seen in Figure 4, with absolute values of Pearson correlation equal to 0.99.

The average diameter contributes to only one of the graphs. Although it was not calculated based on the cell area, using the farthest points of each cell instead, these two measures present a high correlation with Pearson coefficient equal to 0.98. Therefore, tissue groups that present a biologically larger cell area will have larger values of equivalent diameter and diameter. Xylem has a larger cell area when compared to the phloem, so the equivalent diameter and xylem diameter will also be larger as shown in the graphs of Figure 5 A, C and D.

Another relevant conclusion is that all the selected cases involve a measurement referring to the cellular form and another related to density. Figure 5 C, for example, was obtained by pairing entropy 30 and average diameter. This graph presented the largest intercluster distance, but also presented the greatest intracluster distance and (see Table I). Therefore, it was not as effective in the segregation of the groups when compared to the graphs 5 A and B. The graph in Figure 5 D has the entropy attribute 48 on the y-axis and the average equivalent diameter on the x-axis. Finally, the graph in Figure 5 E has the entropy 48 on the y-axis and the average area on the x-axis. It had the smallest intracluster distances as well as the smallest intercluster distances.

The attributes average equivalent diamenter and average area are found on the x-axis in the graphs of the figure 5 A and D and in the graphs of Figure 5 B and E, respectively. The difference between the graphs 5 A and D and the graphs 5 B and E appears on the y axis, which refers to the entropy attribute with sigma 30 and sigma 48. These two attributes distinguish one another only by the sigma value assigned to the Gaussian adopted for the kernel density estimation. In other words, there is a difference in spatial scale between these two measurements, with the entropy 48 characterizing the tissues at a larger scale.

The good segregation between the tissue groups studied was influenced by the discrimination power of the considered paired measures. The high correlation between the measurents of cellular shape and also between those of entropy justify the similar behavior among all presented graphs (see Figure 5).

Table II shows the ratio between the inter and intracluster distances obtained for each pair of tissue in the 5 cases shown in Figure 5. The primary and secondary xylems resulted very close one another in all the considered combinations. Bold values in Table II indicate the separation obtained between the xylems. In the following, we apply the PCA technique to verify if a better separation between the tissues can be obtained.

**TABLE II:** Intercluster distances for the five selected graphs (Figure 5 *A-E*)*. P1 is primary phloem; P2 is secondary phloem, X1 is primary xylem and X1 is secondary xylem*.

|  | A | B | C | D | E |
| --- | --- | --- | --- | --- | --- |
| $P1 \times X1$ | 65,10 | 60,18 | 71,05 | 65,28 | 60,37 |
| $P1 \times P2$ | 68,04 | 67,93 | 68,16 | 63,98 | 63,86 |
| $P1 \times X2$ | 40,21 | 33,75 | 47,02 | 38,28 | 31,42 |
| $X1 \times P2$ | 94,76 | 93,31 | 97,54 | 93,18 | 91,71 |
| $X1 \times X2$ | **33,55** | **35,29** | **32,53** | **31,04** | **32,91** |
| $P2 \times X2$ | 62,94 | 60,73 | 66,03 | 65,28 | 63,15 |

### D. Principal Component Analysis

PCA considers linear combinations of all the original variables and, as a consequence, each new axis will be obtained according to the weighted contribution of each attribute in the original data. Please refer to Section IV C 3 for a more detailed explanation of this method.

Table III shows that the first three main components together account for 74.48% of the original data variance, with the first major component having 47.06%, the second 18.85% and the third 8.02%. In this way, the feature space could be reduced from 33 to only three dimensions, with a relatively small loss of information. Table III also shows the weight of the 33 measures analyzed and, in bold, those that had more relevance in the calculation of the new axes. Interestingly, the only attribute that appeared only in this section was polygonality. The other attributes selected in this section had already provided good separation in the previous analyses.

**TABLE III:**
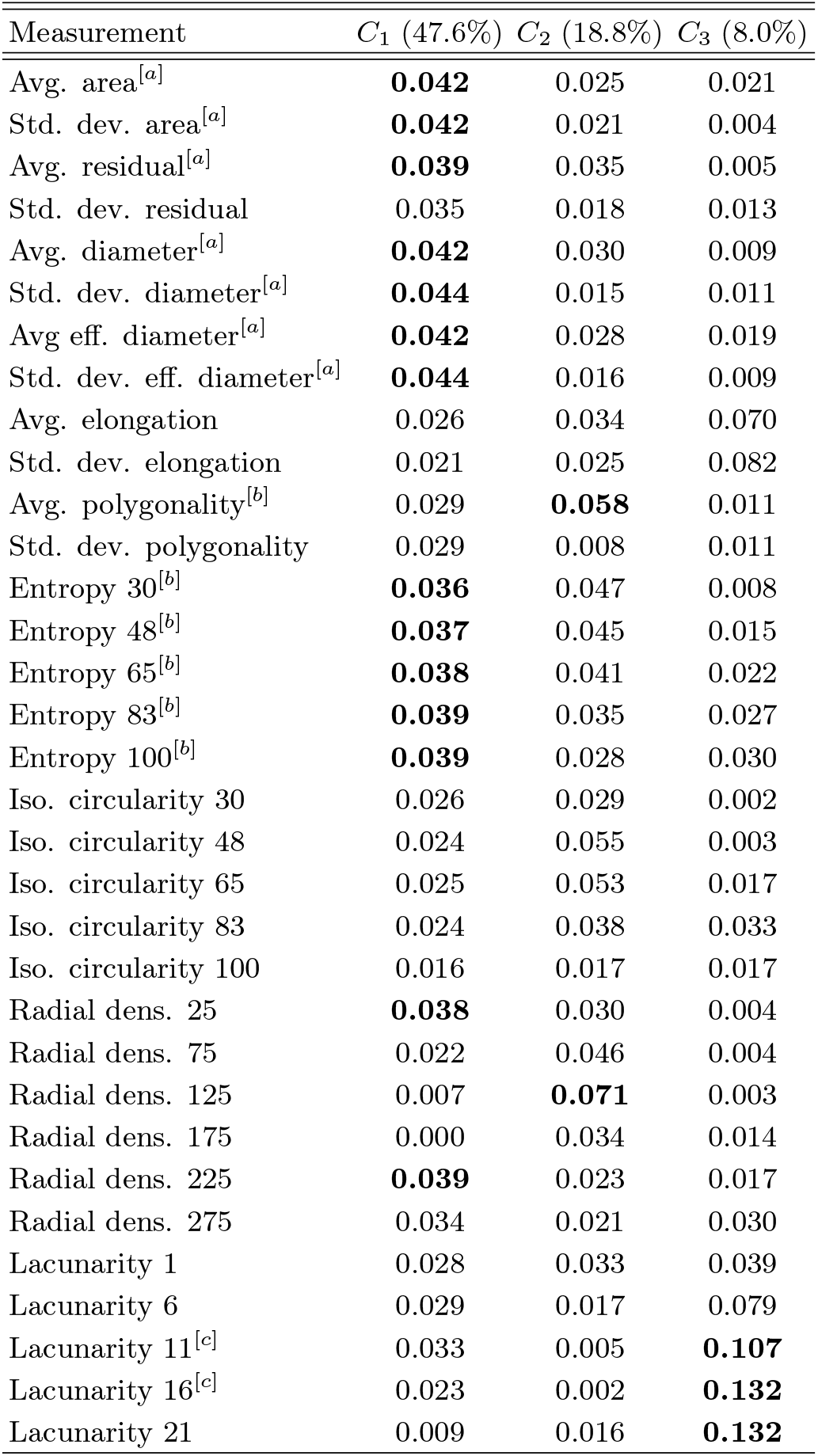
Absolute PCA weights for components *C*_1_*, C*_2_ and *C*_3_. Bold values indicate the largest weights. Measurements in the same connected component in Figure 4 are indicated with a superscript (e.g., for component a).

| Measurement | $C_1$ (47.6%) | $C_2$ (18.8%) | $C_3$ (8.0%) |
| --- | --- | --- | --- |
| Avg. area<sup>[a]</sup> | **0.042** | 0.025 | 0.021 |
| Std. dev. area<sup>[a]</sup> | **0.042** | 0.021 | 0.004 |
| Avg. residual<sup>[a]</sup> | **0.039** | 0.035 | 0.005 |
| Std. dev. residual | 0.035 | 0.018 | 0.013 |
| Avg. diameter<sup>[a]</sup> | **0.042** | 0.030 | 0.009 |
| Std. dev. diameter<sup>[a]</sup> | **0.044** | 0.015 | 0.011 |
| Avg eff. diameter<sup>[a]</sup> | **0.042** | 0.028 | 0.019 |
| Std. dev. eff. diameter<sup>[a]</sup> | **0.044** | 0.016 | 0.009 |
| Avg. elongation | 0.026 | 0.034 | 0.070 |
| Std. dev. elongation | 0.021 | 0.025 | 0.082 |
| Avg. polygonality<sup>[b]</sup> | 0.029 | **0.058** | 0.011 |
| Std. dev. polygonality | 0.029 | 0.008 | 0.011 |
| Entropy 30<sup>[b]</sup> | **0.036** | 0.047 | 0.008 |
| Entropy 48<sup>[b]</sup> | **0.037** | 0.045 | 0.015 |
| Entropy 65<sup>[b]</sup> | **0.038** | 0.041 | 0.022 |
| Entropy 83<sup>[b]</sup> | **0.039** | 0.035 | 0.027 |
| Entropy 100<sup>[b]</sup> | **0.039** | 0.028 | 0.030 |
| Iso. circularity 30 | 0.026 | 0.029 | 0.002 |
| Iso. circularity 48 | 0.024 | 0.055 | 0.003 |
| Iso. circularity 65 | 0.025 | 0.053 | 0.017 |
| Iso. circularity 83 | 0.024 | 0.038 | 0.033 |
| Iso. circularity 100 | 0.016 | 0.017 | 0.017 |
| Radial dens. 25 | **0.038** | 0.030 | 0.004 |
| Radial dens. 75 | 0.022 | 0.046 | 0.004 |
| Radial dens. 125 | 0.007 | **0.071** | 0.003 |
| Radial dens. 175 | 0.000 | 0.034 | 0.014 |
| Radial dens. 225 | **0.039** | 0.023 | 0.017 |
| Radial dens. 275 | 0.034 | 0.021 | 0.030 |
| Lacunarity 1 | 0.028 | 0.033 | 0.039 |
| Lacunarity 6 | 0.029 | 0.017 | 0.079 |
| Lacunarity 11<sup>[c]</sup> | 0.033 | 0.005 | **0.107** |
| Lacunarity 16<sup>[c]</sup> | 0.023 | 0.002 | **0.132** |
| Lacunarity 21 | 0.009 | 0.016 | **0.132** |

Figure 6 illustrates the scatterplot obtained through the PCA projection. In Figure 6 A we have the first principal component (*C*_1_) on the x-axis and the second (*C*_2_) on the y-axis. Together they account for 66.45% of the total variance of the original data. In this graph, there is a small region of overlap between the clusters for the primary and the secondary xylem. In Figure 6 B, *C*_1_ is the x-axis while the third principal component (*C*_3_) is the y-axis. Together, the two axes accumulate 55.63 % of the total variance of the original data. In this graph, the primary and secondary xylems are almost overlapping.

**FIG. 6:**
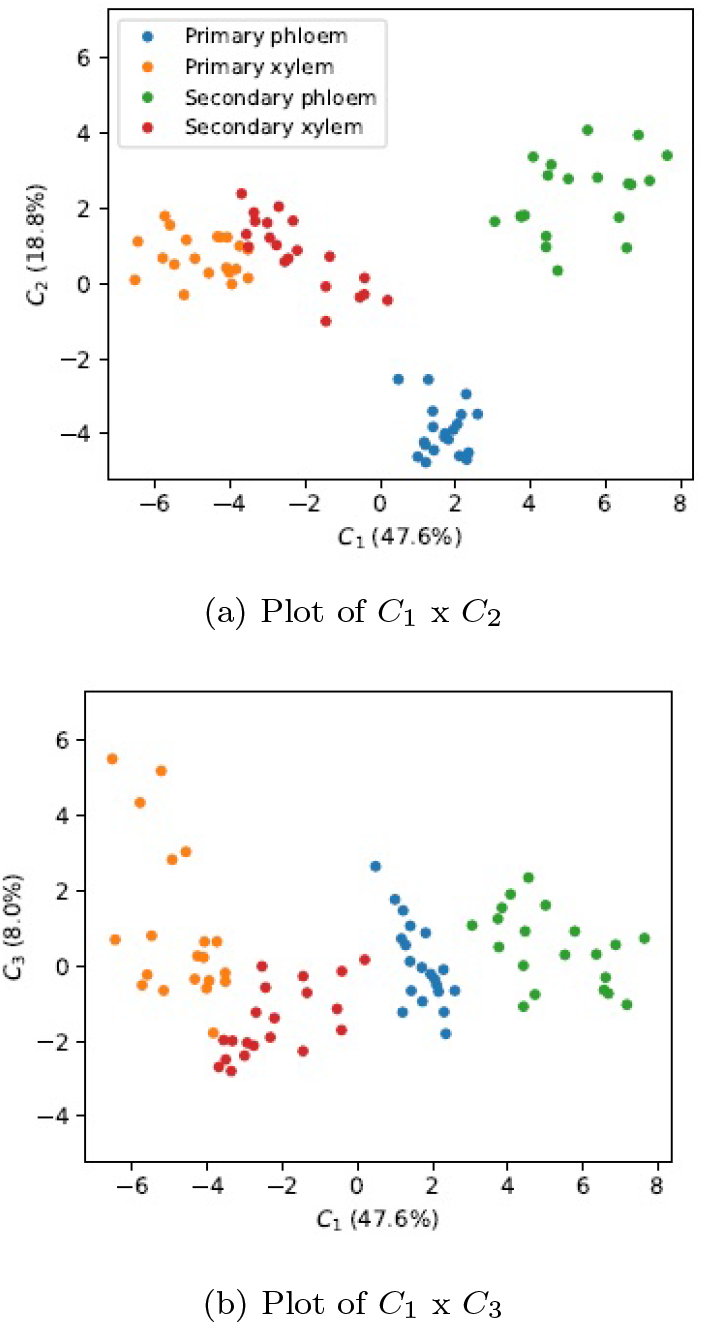
Scatterplot obtained through the PCA projection.

The intercluster distances of the graphs in Figure 6 are presented in Table IV. The results show that Figure 6 B is composed of clusters more dispersed along the axes when compared to Figure 6 A, since the intracluster distance is larger for this graph.

**TABLE IV:** Interclusters and intraclusters average distance and relation between distances (*r*) for the PCA projection. *C*_1_ *is first principal component and C*_2_ *is second principal component*.

| | Inter | Intra | $r$ |
| --- | --- | --- | --- |
| $C_1 \times C_2$ | 58,46 | 13,11 | 4,46 |
| $C_1 \times C_3$ | 44,71 | 18,76 | 2,38 |

Although the graph obtained for *C*_1_ × *C*_3_, see Figure 6 B, has presented a greater distance between the primary xylem (*X*1) and the secondary xylem (*X*2) if compared to *C*_1_ × *C*_2_, see Figure 6 B, the distance between all other groups has decreased, as evidenced in the Table V. The greater proximity between the groups may generate errors in a classification system. Thus, among the results obtained by the PCA, Figure 6 A presented the best separation.

**TABLE V:** Intercluster distances for the PCA projection (6 *A and B*)*. P1 is primary phloem; P2 is secondary phloem, X1 is primary xylem and X1 is secondary xylem*.

|  | A | B |
| --- | --- | --- |
| $P1 \times X1$ | 69,24 | 46,65 |
| $P1 \times P2$ | 75,06 | 25,80 |
| $P1 \times X2$ | 60,17 | 32,93 |
| $X1 \times P2$ | 72,95 | 70,84 |
| $X1 \times X2$ | 18,04 | 34,42 |
| $P2 \times X2$ | 55,31 | 57,64 |

## III. DISCUSSION

Recent advances in image analysis and pattern recognition have paved the way to several applications in biology. In particular, we have the interesting problem of characterizing plants images so as to identify structural elements such as tissues and cells. Results from this type of research would contribute to developing methods for content-based retrieval, diagnostics, species identification, among other possibilities. The current work addressed the characterization of primary and secondary vascular bundles by using several methods and concepts from image analysis and artificial intelligence, with special attention given to the issue of finding a reduced set of measurements capable of good separation of the considered types of tissues.

A total of 20 images containing primary and secondary vascular bundles of the stem of *Dracaena marginata* was analyzed. Although these tissues have the functionality of transporting water and nutrients, each of them presents particularities and is involved in specific subfunctions, resulting in unique characteristics making them biologically and computationally differentiable.

A set of 33 morphological, density and regularity measurements was extracted from each image, which was then studied using three approaches: i) individual analysis; ii) pairwise analysis and iii) PCA projection. The aim was to search for the attributes that could segregate well the primary and secondary vascular bundles of monocotyledon with secondary growth. This work showed that the most relevant attributes always belonged to the subgroup of attributes: area, diameter, entropy, polygonality, lacunarity and radial density, regardless of the approach used (individual analysis, pairwise analysis and PCA). In addition, the results for pairwise analysis resulted in better results than the other two approaches used in this work.

The pairwise measurement analysis was able to completely separate the vascular tissue groups of the *Dracaena marginata*. More specifically, a combination between the entropy 30 and average equivalent diameter measurements led to the best separation between tissues and a good distance between the groups. In general, the entropy with sigma of 30 or 48 when analyzed together with measures concerning the cellular form, such as area and diameter, also presented satisfactory results for tissue segregation.

Biologically, the entropy, area and diameter characteristics were expected to play an important role in distinguishing vascular plant tissues. The entropy can be interpreted as the degree of regularity and organization of a geometric structure such as the considered tissues. More organized tissues have smaller entropy. The primary tissues are organized according to the axial system while the secondary tissues are organized according to the axial and radial systems (Cutler et al., 2009; Raven et al., 2014). Thus, the secondary tissues are more regularly organized and, therefore, have lower values of entropy when compared to the primary counterparts, as presented in the sections above. The difference in entropy between xylem and phloem may be related to the condition of the constituent cells. The phloem is composed of living cells, which can lead to a less organized tissue when compared to the phloem, which is composed of dead and lignified cells.

The results related to the cellular shape led to a relevant biological interpretation. Xylem is the tissue responsible for transporting water and minerals, while the phloem carries the organic compounds resulting from photosynthesis. Thus, the transport capacity of the xylem exceeds that of the phloem since the fluid transport rate is higher in the xylem than in the phloem (Hölttä et al., 2009; Wu et al., 2011; Lucas et al., 2013). This hydraulic difference between the tissues justifies the distinction between the diameter of the xylem and the phloem (Santarosa et al.; 2016). The xylem, because it has a greater flow capacity, tends to consist of larger cells when compared to the phloem (Jura-Morawiec; 2015). The area and diameter resulted in a high and positive Pearson correlation coefficient, indicating that cells with large areas also tend to have a high diameter. Therefore, all measures related to the cellular shape considered in the analysis showed a similar behavior when analyzed together with the entropy.

The other attributes highlighted in this work (lacunarity, polygonality and radial density) also have biological relevance. The lacunarity characterizes the homogeneity of the distribution of holes in an image. If the image contains irregularly distributed holes of many different sizes, the lacunarity of the image is high. Contrariwise, if the holes have similar size and are homogeneously distributed, the lacunarity is low (Rodrigues et al., 2005). The main constituent cells of xylem of monocot with secondary growth are tracheids (Raven et al., 2014). These cells, during their development, undergo a process called intrusive growth (Jura-Morawiec, 2017), that is, a cell will penetrate between the adjacent cells (Cutler et al., 2009). According to de Oliveira Santos and Nogueira (1977), intrusive growth occurs in both the primary and secondary tissues, but in the latter it is more significant. This process results in the occupation of intercelular spaces (Wenham and Cusick, 1975; Jura-Morawiec, 2017), thus, it may promote a greater homogeneity in the distribution of the secondary xylem in relation to the primary one. The intrusive growth does not occur in the phloem because it is a characteristic of tracheid cells.

The polygonality is the measurement that quantifies the regularity of the angular distribution of neighboring cells. The phloem constituent cells are alive throughout the plant development, while the xylem cells, as they mature, undergo lysis of the cytoplasm and parts of the cells wall give rise the formation of a secondary lignified wall culminating in cell death, which then become apt for transport processes. As the xylem is composed of mostly of dead cells, the changes that occur in the xylem are irreversible (Raven et al., 2014). This characteristic of this tissue indicate that in a tissue composed mostly of living cells, it is reasonable to expect that the properties of neighboring cells have a greater variation than in dead tissues (Jura-Morawiec, 2017). In relation the origin of tissues: primary vascular tissues reach their maximum degree of differentiation and remain functional throughout the individual’s life without further differentiation (Raven et al., 2014). Thus, it is reasonable to expect that primary tissues exhibit less variation in the angular distribution of neighboring cells when compared to secondary tissues that undergo differentiation throughout plant life.

The radial density is calculated from the center of mass of the centroid of each cell of the tissue. The tissue organization itself of the vascular bundles justifies the relevance of this attribute. As the vascular bundle type studied was amphivasal, that is, concentric with the phloem being surrounded by the xylem (Tomlinson and Zimmermann, 1969), this attribute had interesting results mainly in the characterization of the primary phloem.

The reported results suggest that image analysis methods can help the classification process of vascular bundles, since clusters referring to each tissue type were well segregated in a *2D* space. The obtained results pave the way to several applications such as in content-based retrieval, species identification and diagnosis. Other future works include extending the proposed methodology to other plant species, as well as other types of vascular bundles. It would also be interesting to study the evolution of these structures along time and the position along the stem.

## IV. EXPERIMENTAL PROCEDURES

### A. Image Database

The image database was generated using slides with transverse sections of the stem of *Dracaena marginata*, which was kindly supplied by Prof. Dr. Marcos Arduin of the Department of Botany at Federal University of São Carlos (UFSCar).

Image acquisition was carried out by a photo camera (Canon - PowerShot A650IS 12.1MP Digital Camera) attached to an optical microscope (Carl Zeiss - Axiostar Plus Transmitted Light Microscope). All images were captured with 2400x magnification. Images of the primary and secondary vascular bundles were captured and separated into two groups according to the bundles specification, that is, primary or secondary tissue. The groups consisted of 20 images that were analyzed separately as described in Section IV B.

### B. Image Analysis

#### 1. Cell segmentation

The primary and secondary vascular bundles are composed of xylem and phloem, so this step was performed with the intention of manually segmenting these tissues into the two types of bundles studied. Segmentation was done using the software *Paint.net* (*v3.08*) [dotPDN LLC] according to the following steps:

1. the original image was used as background for segmentation;
2. a transparent mask was added over the original image;
3. xylem cells were outlined and filled with gray levels from 1 to 255;
4. the unselected region, that is, the region of the image that did not correspond to xylem cells was filled with the level 0 gray tone.
5. the mask containing the cells was saved;
6. on the original image, a new transparent mask was added;
7. the cells related to the phloem were outlined and filled with gray levels from 1 to 255;
8. the unselected region, that is, the region of the image that did not correspond to phloem cells was filled with gray level 0.

The steps were repeated for all images of both the primary and secondary vascular bundle groups. At the end of this stage, the group consisting of images of primary vascular bundles was subdivided into xylem and primary phloem. Similarly, the group consisting of images of secondary vascular bundles was subdivided into secondary xylem and phloem. Figure 7 illustrates the separation of the groups that will be considered throughout this study.

**FIG. 7:**
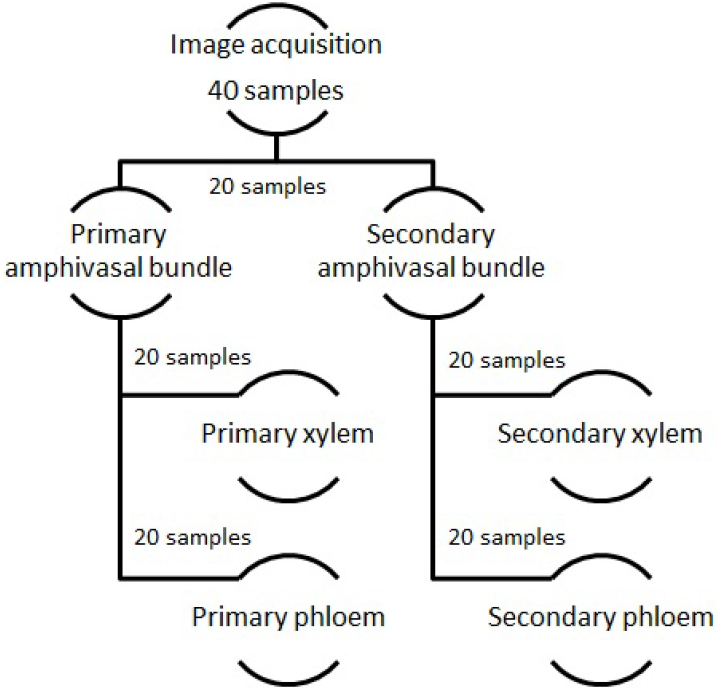
Division of four groups: primary xylem and phloem, secondary xylem and phloem. Each group is composed of 20 tissue samples.

#### 2. Measurements related to cell shape

Measurements described in this section were calculated for individual cells, and the average and standard deviation over all cells in a tissue was obtained for tissue characterization.

**Cell area:** The number of pixels representing the cell (da Fontoura Costa and Cesar Jr, 2009).

**Convex hull residual:** First, the convex hull (De Berg et al., 2000) of the cell is calculated. Then, the convex hull residual is given by the difference between the convex hull area and the cell area (da Fontoura Costa and Cesar Jr, 2009).

**Max cell distance:** The maximum distance between any two points in the cell (da Fontoura Costa and Cesar Jr, 2009).

**Equivalent diameter:** The equivalent diameter is calculated as

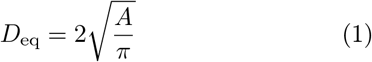

where *A* is the cell area. *D*_eq_ is the diameter of a disk having the same area as the cell (da Fontoura Costa and Cesar Jr, 2009).

**Elongation:** Let 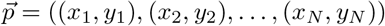 be a vector containing the image row and column positions of all *N* pixels representing the cell. The principal component analysis (PCA) (Jolliffe, 1986) is applied to vector 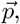 resulting into two axes that can represent the shape with optimum variance. The square root of the ratio between the eigenvalues of the first and second principal components indicates the overall elongation of the shape (da Fontoura Costa and Cesar Jr, 2009).

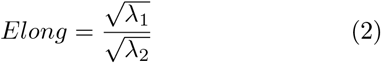

where λ_1_ e λ_2_ are eigenvalues of the first and second principal components

#### 3. Density measurements

The density measurements are based on the distribution of the cell centroids. The entropy and isoline circularity measurements (described below) were calculated over an unsupervised estimation of the probability distribution of cell centroids. This is done by applying a Gaussian kernel density estimation methodology (Duda et al., 2012), where each centroid is treated as an unitary impulse. The sigma parameter of the kernel Gaussian function was varied between the following values: 30, 48, 65, 83 and 100. Larger sigma values lead to smoother probability distributions and, therefore, to measurements that quantifies the tissues at larger spatial scales.

**Entropy of centroid density:** The Shannon (Shannon, 1948) entropy of the estimated probability distribution.

**Isoline circularity:** By thresholding the probability distribution at the 0.95th percentile, a binary image is obtained. Then, the circularity of this binary image is calculated (da Fontoura Costa and Cesar Jr, 2009). This measurement is calculated as

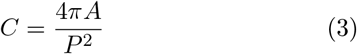

where *A* and *P* are, respectively, the area and perimeter of the thresholded image.

**Centroid radial distribution:** First, the centroid c of the set of all cells in the image is calculated. Then, an annulus centered on *c* and having inner radius *r_i_* and outer radius *r_i_* + ∆r is defined. Next, the density of cell centroids contained in the annulus is obtained. This is calculated as the ratio between the number of cell centroids inside the annulus and the area of the annulus. A descriptor 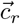 can be defined, containing the cell centroid density for distinct annulus radii values. In the analysis we consider *r_i_* = *i* * ∆r for all *i* in the range [0, 5], and ∆*r* = 50 pixels.

#### 4. Structural regularity measurements

The measurements in this section are related to the regularity of the cells distribution.

**Lacunarity:** This property characterizes the homogeneity of the distribution of empty spaces (holes) in an image (Rodrigues et al., 2005). If the image contains irregularly distributed holes of many different sizes, the lacunarity of the image is high. Contrariwise, if the holes have similar size and are homogeneously distributed, the lacunarity is low. The "holes" in our tissue image were generated by taking the negative of the binary mask containing the manually segmented cells. That is, each cell in the original image becomes a hole in the negative image. The lacunarity was calculated within a region bounded by a radius R around a given pixel of the image. The parameter R ranged from the following values: 1, 6, 11, 16 and 21.

**Polygonality:** The cell centroids are used for defining a respective Delaunay triangulation (De Berg et al., 2000). For a given cell *i*, the Delaunay triangulation is used for defining the neighboring cells, Γ*_i_*, of this cell. These cells tend to be spatially close to cell *i*. Let’s define a line *l_ij_* going from centroid *i* to centroid *j*, where cell *j* belongs to the neighborhood Γ*_i_* of cell *i* and *j* = 1, 2 …, *N_i_*. *N_i_* being the number of neighbors of cell *i*. Respective angles *θ_j,j_*_+1_ between each consecutive line (in a clockwise direction) are then defined (da Fontoura Costa et al., 2006). The polygonality is then defined as

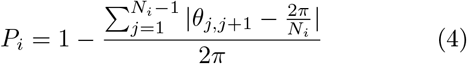

This property quantifies the regularity of the angular distribution of neighboring cells, that is, the similarity between the values *θ_j,j_*_+1_ and the angle 2*π*/*N_i_*. If *θ_j,j_*_+1_ = 2*π*/*N_i_* for all *j*, the neighbors of cell *i* have a regular angular distance.

### C. Pattern Recognition

#### 1. Individual Measurement Analysis

Let *X* be a random variable, described in terms of its probability density function (*f*_(_*_x_*_)_). So, for any values *a* and *b*, the probability that X is comprised between *a* and *b* is given as:

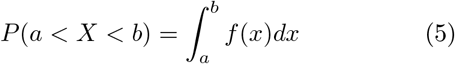

for values of *a* < *b*. The probability density function is estimated by convolving the histogram of variable *x* with a Gaussian kernel (Duda et al., 2012). In this work, the probability denstity function was plotted using the *histfit* function in the software *Matlab*.

The graphs reveal the potential of each measurement for separating the distinct tissue types considered. According to Bayesian inference, the smaller the overlap region between two curves, the better the groups will be separated and, consequently, the classes will be better defined. Therefore, the graphs were analyzed and those leading to the best discrimination between the tissue types were selected.

#### 2. Pairwise Measurement Analysis

Generally, a single attribute is not able to extract enough information for a good tissues characterization, so in this step we considered pairs of measurements. This approach is justified because, when working with two attributes, we increase the number of dimensions to two (*2D*) and this may promote a better separation of the groups.

Often, it is important to avoid pairs of characteristics that have a high degree of correlation, as they may have great redundancy. An analysis of Pearson’s correlation coefficient (*p*) (Pearson, 1896) was performed in order to quantify the degree of correlation between two variables and to verify if these correlations are positive or negative. The Pearson correlation between two random variables *X* and *Y* can be calculated as

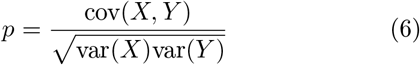

in which cov(*X*, *Y*) corresponds to the covariance between the variables *X*, and *Y* and var(*X*) is the variance of variable *X*.

Values of *p* such that *p* ≈ 1 or *p* ≈ − 1 indicate that the variables analyzed are strongly correlated and, therefore, may be redundant.

Scatterplots between each pair of measurements were also obtained. In order to study the dispersion between and within tissue groups, we used the intercluster and intracluster distance matrices, respectively (Duda et al., 2012). For the calculation of both analyzes, the variables were normalized between 0 and 1.

The intercluster distance matrix was calculated taking into account the euclidean distance of the centroids from each tissue cluster. The centroid 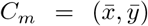 was estimated as the mean of the coordinate values of the points, as described below.

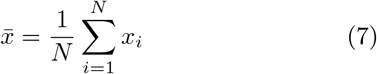

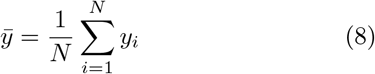

where *N* is the number of points in the category.

The Euclidean distance was calculated as described below:

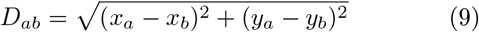

where, *x_a_* and *x_b_* are the centroids of the coordinate x between two categories and *y_a_* and *y_b_* are the centroids of the coordinate x between two categories.

The intracluster distance was calculated as the average euclidean distance between all points of the same tissue. Let i and j be two points of the same category, the Euclidean distance between i and j was calculated as follows:

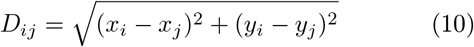

All distance values were summed and, at the end, the arithmetic mean was calculated.

The separation between the groups can, then, be quantified by dividing the intercluster distance by the intracluster distance (r).

#### 3. Principal Component Analysis

Principal Component Analysis (*PCA*) (Jolliffe, 1986) is a technique used for reducing the dimensionality of a dataset while preserving as much as possible the variance of the original data, thus eliminating redundancies in the data (Vasconcelos, 2007; Varella, 2008). This analysis can be understood as a linear transformation of the original variables (Vasconcelos, 2007) onto a new space defined by set of uncorrelated measurements. A possible procedure to calculate PCA is presented in Figure 8.

**FIG. 8:**
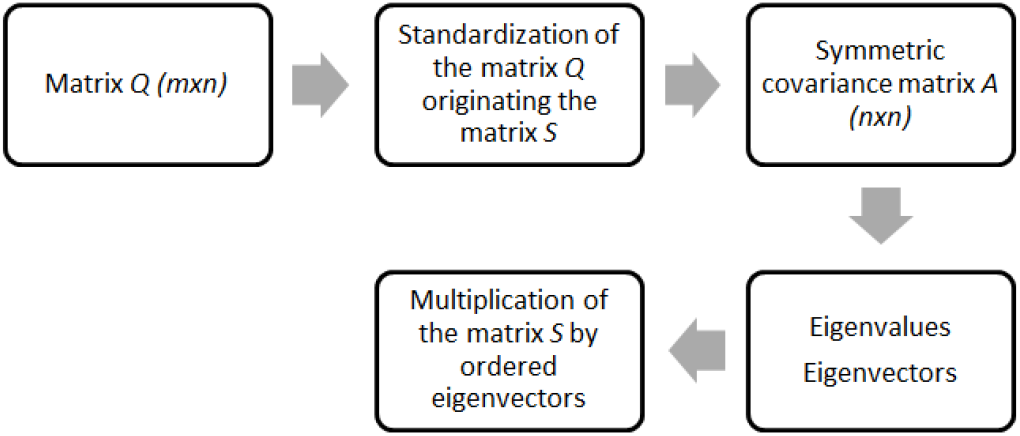
Flowchart illustrating the summarized steps to perform the principal component analysis.

After calculating a set of *n* measurements for each image, the obtained values are organized as a data matrix (Q), having size *m* × *n*. Henceforth, *m* is the number of samples (images) and *n* the number of measurements. As the units of the measurements can vary in magnitude between the attributes, it is convenient to standardize the matrix *Q*. The standardization is done following the equation below, yielding the matrix *S*.

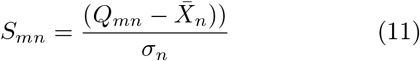

in which 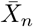 is the average and *σ_n_* the standard deviation of measurement *n*.

The covariance among each column of *S* is then calculated, resulting in the covariance matrix *A*, having size *n* × *n*. The eigenvalues and eigenvectors of matrix *A* are then calculated (Varella, 2008). The *L* principal components correspond to the eigenvectors associated to the *L*-largest eigenvalues of *A*.

After the determination of the main components, the contribution of each main component to the data projection can be calculated. This contribution is given in percentage and represents the ratio of total variance explained by the main component. Usually, principal components accumulating approximately 70% or more of the total variance ratio are chosen (Varella, 2008). Finally, the matrix *S* is multiplied by the selected eigenvectors, defining new, uncorrelated, values to each tissue image.

## Acknowledgments

This study was financed in part by the Coordenação de Aperfeiçoamento de Pessoal de Nível Superior - Brasil (CAPES) - Finance Code 001. Moreover, Cesar H. Comin thanks FAPESP (grant no. 2018/09125-4) for financial support. Luciano da F. Costa thanks CNPq (grant no. 307333/2013-2) and NAP-PRP-USP for sponsorship. This work has been supported also by the FAPESP grant 2015/22308-2.

We also thank Professor Marcos Arduin of the Department of Botany at Federal University of São Carlos (UFSCar) for the biological material used in this study.

